# Transketolase of *Staphylococcus aureus* is involved in the control of master regulators of stress response during infection

**DOI:** 10.1101/538900

**Authors:** Xin Tan, Elodie Ramond, Anne Jamet, Baptiste Decaux-Tramoni, Marion Dupuis, Daniel Euphrasie, Fabiola Tros, Ivan Nemazanyy, Jason Ziveri, Xavier Nassif, Alain Charbit, Mathieu Coureuil

## Abstract

*Staphylococcus aureus* is a leading cause of both acute and chronic infections in humans. The importance of the pentose phosphate pathway (PPP) during *S. aureus* infection is currently largely unexplored. Here, we focused on one key PPP enzyme, transketolase. We showed that inactivation of the unique gene encoding transketolase activity in *S. aureus* USA300 (∆*tkt*) led to drastic metabolomic changes. Using time-lapse video imaging and mice infection, we observed a major defect of the ∆*tkt* strain compared to wild-type strain in early intracellular proliferation and in the ability to colonise kidneys. Transcriptional activity of the two master regulators Sigma B and RpiRc was drastically reduced in the ∆*tkt* mutant during host cells invasion. The concomitant increased RNAIII transcription, suggests that TKT-or a functional PPP-strongly influences the ability of *S. aureus* to proliferate within host cells by modulating key transcriptional regulators.

## Background

*Staphylococcus aureus* is a leading cause of severe skin and soft tissue infections, pneumonia, endocarditis and bone and joint infections [1, 2]. *S. aureus* invade and survive within eukaryotic host cells [3-9]. This is likely to play an important role in chronicity and treatment failures [10-12]. To evade host defences and silently persists within human cells [13, 14], *S. aureus* can dynamically switch phenotypes from a highly aggressive and cytotoxic phenotype to a metabolically inactive phenotype associated with decrease TCA activity [15-18]. Reprogramming of the TCA cycle notably involves the glycolysis/gluconeogenesis and pentose phosphate pathway (PPP) [19].

The contribution of glycolysis/gluconeogenesis to intracellular persistence of *S. aureus* has been recently studied [20-23]. In contrast, the precise role of the PPP is unexplored. The PPP is important for producing nucleotide precursors for DNA synthesis and repair as well as precursors for aromatic amino acids or peptidoglycan synthesis. In addition, the oxidative branch of the PPP contributes to bacterial tolerance to oxidative stress by generating the redox molecule NADPH,H^+^, which is the substrate for other reducing agents such as glutathione. The activity of the PPP has been shown to be frequently increased in Gram-positive pathogens in response to environmental stresses and in infection models [24-26] or in some hospital-acquired isolates [27]. While it has been suggested that the transcriptional regulator RpiRc might control the PPP [28-30], no gene regulation has been yet associated with any change in its activity in *S. aureus*.

Here, we study the role of PPP during the intracellular persistence of *S. aureus* by focusing on one of the key enzyme of this pathway, transketolase (TKT). *tkt* genes are found in a wide range of organisms including bacteria, plants and mammals, underlining the ubiquitous role of this enzyme in biology. We show here that *tkt* inactivation in *S. aureus* USA300 leads to dysregulation of whole cell metabolism. This correlates with impaired proliferation of intracellular staphylococci *in vitro* and kidney colonisation defect in infected mice. Unexpectedly, we found that the *tkt* defective mutant was unable to regulate the two master regulators of virulence RpirC and SigB, leading to a dramatic increase of RNAIII during intracellular invasion of human cells. Altogether our data suggest a major role of the PPP in the control of *S. aureus* virulence.

## Methods

### Strains, Cells, Culture conditions and Infection

*S. aureus* USA300-LAC (designated USA300-WT) was provided by the Biodefense and Emerging Infections Research Resources (BEI). The GFP-expressing strain, the Δ*tkt* mutant strain and the complemented strain were constructed and cultured as described in **supplementary methods**. Strains and plasmids used are summarized in **supplementary table S1A**.

*S. aureus* were grown in brain heart infusion (BHI) or chemically defined medium (CDM) supplemented with ribose or fructose or glycerol when it was required [31].

EA.hy926 endothelial cells (ATCC CRL-2922) were grown in DMEM 10 % FBS (5 % CO_2_ at 37 °C). When indicated cells were incubated with no glucose DMEM supplemented with glucose, ribose or fructose at a concentration of 25mM. EA.hy926 expressing nuclear mKate2 were transduced with the IncuCyte^®^ NucLight Red Lentivirus.

Cells were infected at a multiplicity of infection (MOI) of 1 for 1 h and washed three times with 100 µL of phosphate buffer saline (PBS) containing 300 µg/mL gentamicin to remove extracellular bacteria. Cells were then incubated in PBS-gentamicin (50µg/ml). Gentamicin is an antibiotic with a high bactericidal effect on *S. aureus* USA300 (CMI=2 µg.mL^-1^) and a very poor penetration inside eukaryotic cells. Its use was necessary to abolish extracellular proliferation.

LAMP-1 colocalization assay in EA.hy926 infected cells is described in the **supplementary methods**.

### Time lapse microscopy

EA.hy926 cells expressing nuclear mKate2 were seeded in ImageLock 96-well plates (Essen BioScience Inc). Cells were infected with WT and Δ*tkt* mutant of the GFP-expressing USA300 strain. Plates were incubated and monitored at 5% CO2 and 37°C for 24 hours in the fully automated microscope Incucyte^®^ S3 (Essen BioScience). The detailed analysis is further described in the **supplementary methods**.

### Transcriptional analyses

The bacteria were recovered from lysed cells or planktonic overnight culture. Nucleic acids were released by resuspending bacteria in TE buffer containing lysostaphin. RNAs recovery, Reverse transcription and PCR were performed as described in the **supplementary methods**.

### Metabolomic analyses

Metabolite profiling of *S. aureus* isolates grown to stationary phase in BHI Broth was performed by liquid chromatography–mass spectrometry (LC-MS) as described in the **supplementary methods** [9, 32, 33].

### Transketolase activity assay

TKT activity was analysed on bacteria grown to stationary phase in BHI as described in the **supplementary methods** [34]. TKT activity was expressed as units per microgram of total protein. One unit of enzyme was defined as the amount of enzyme that oxidized 1 µmol of NADH per minute.

### *In vivo* infection

Mice were infected intravenously (iv) in the tail vein as described in the **supplementary methods**. For bacterial burden determination, mice were euthanized after 4, 24, 48 and 72 hours. Sequential dilutions of homogenized preparations from spleen and kidney were spread onto TS agar plates.

### Statistics

Statistical significance was assessed using one-way analysis of variance (ANOVA) with Dunnett’s correction, unpaired two-tail Student’s t-test, or Kruskal Wallis test. P values of *p*<0.05 were considered to indicate statistical significance.

### Ethics statement

All experimental procedures involving animals were conducted in accordance with guidelines established by the French and European regulations for the care and use of laboratory animals (Decree 87–848, 2001–464, 2001–486 and 2001–131 and European Directive 2010/63/UE) and approved by the INSERM Ethics Committee (Authorization Number: 75-906).

## Results

### Transketolase inactivation in *S. aureus* USA300

*S. aureus* genomes possess a unique transketolase-encoding gene, designated *tkt,* which is highly conserved within the species. Transketolase (TKT) is an enzyme of the non-oxidative branch of the PPP involved in two main reversible enzymatic reactions: i) Fructose-6-P + Glyceraldehyde-3-P <-> Erythrose-4-P + Xylulose-5-P; and ii) Sedoheptulose-7-P + Glyceraldehyde-3-P <-> Ribose-5-P + Xylulose-5-P (**Supplementary Figure S2A)**. Several TKT structures have been solved [35] and allow the modelisation of the *S. aureus* TKT **(Supplementary Figure S1B).** The protein of *S. aureus* is 662 amino acid long and harbours three domains **(Supplementary Figure S1A)**. Of note, although ubiquitously expressed in Eukaryotes and Bacteria, *S. aureus* and *Homo sapiens* TKT proteins share only 22.4% amino acid identity **(Supplementary Figure S1C)**. We constructed a chromosomal deletion of the *tkt* gene in *S. aureus* strain USA300 (∆*tkt* strain), and generated a complemented strain (designated Cp-*tkt*) by expressing the wild-type *tkt* allele preceded by its own promoter in the ∆*tkt* strain. Gene inactivation was confirmed by quantification of the transketolase activity (**Supplementary Figure S2)**.

The ∆*tkt* mutant showed only a slight growth decrease compared to the WT strain in complete BHI Broth or in Dulbecco's Modified Eagle Medium (DMEM) (**Figure 1A**). In contrast, the ∆*tkt* mutant strain was unable to grow in chemically defined medium [31] supplemented with ribose (which can be converted to Ribose-5-phospate by ribokinase) (**Supplementary Figure S2C**). In all assays, functional complementation was observed in the Cp-*tkt* strain.

**Figure 1.**
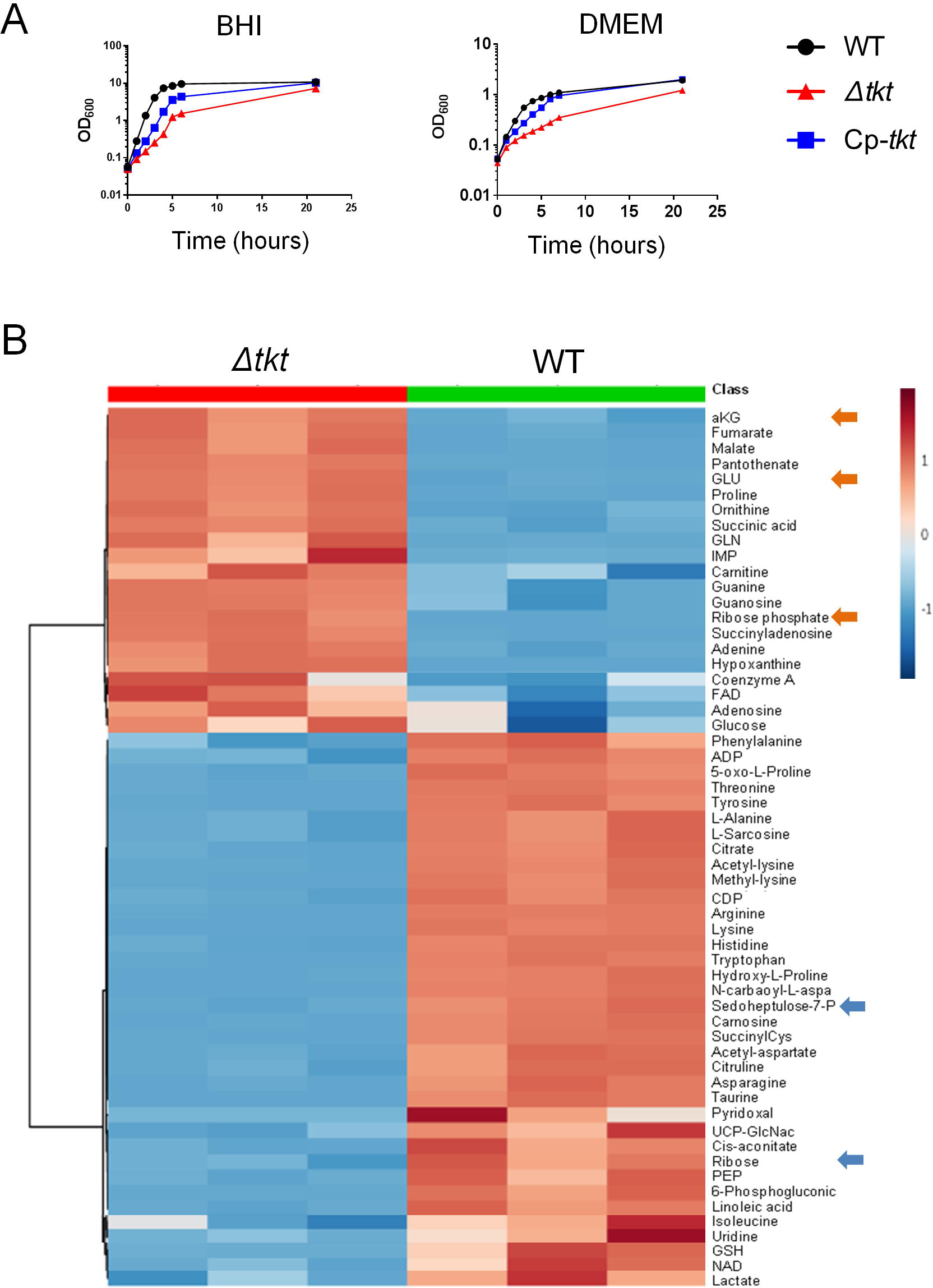
Phenotypic characterization of the ∆*tkt* mutant. **A. Growth of USA300 wild type (WT), Δ*tkt* mutant (Δ*tkt*) and its complemented derivative (Cp-*tkt*).** USA300 WT, ∆*tkt* and Cp-*tkt* strains were grown in BHI Broth or DMEM for 21 hours at 37°C. Optic densities were measured every-hour from 0 to 8 hours and then at 21 hours. Each panel is a representative experiment. **B. Metabolomic analysis of the ∆*tkt* mutant strain.** Quantitative metabolomic analysis was performed by ion chromatography and tandem mass spectrometry (IC-MS/MS) after overnight cultures, in BHI Broth, from WT USA300 and ∆*tkt* mutant strains (see **Table S1**). The heatmap, obtained using MetaboAnalyst (https://www.metaboanalyst.ca), illustrates the metabolic shift between the two strains.

We next performed a whole cell metabolomic analysis on WT and ∆*tkt* strains grown for 24h in BHI Broth (*i.e.,* when identical numbers of cells were recorded) to evaluate the global impact of *tkt* deletion on bacterial metabolism (**Figure 1B and Supplementary Figure S3 and Table S1**). Remarkably, we observed a massive accumulation of ribose-5-phosphate (R5P; 56-fold increase) concomitant with a decrease in ribose content (4-fold decrease) in the ∆*tkt* mutant compared to WT. This accumulation was associated with an increased amount of metabolites derived from Ribose-5P, such as inosine-5-monophosphate (IMP), xanthosine-5-monophosphate (XMP) and hypoxanthine. We also observed a 2.5-fold decrease in D-sedo-heptulose-7P amount in the ∆*tkt* mutant compared to WT accompanied by a significant decrease in tyrosine, histidine and tryptophan amounts (5, 10 and 12 fold respectively), three aromatic amino-acids produced from the PPP. The glutamate pathway seemed to be significantly more active in the ∆*tkt* strain compared to WT as the amounts of glutamate, proline and ornithine were all increased. The higher amounts of glutamate, possibly fuelling the TCA cycle at the level of α-ketoglutarate, could be responsible for the increased expression of some TCA cycle metabolites (*i.e.,* fumarate, malate, succinate and α-ketoglutarate) in the ∆*tkt* strain. Of note, ATP content was not altered in ∆*tkt* strain.

These results demonstrate that deregulation of the PPP by *tkt* inactivation has direct effect on the whole cell metabolism, including aromatic amino acid production, TCA activity and pathways involved in adenine, guanine or peptidoglycan synthesis.

### *tkt* is essential for intracellular proliferation of *S. aureus*

Intracellular staphylococci exist as two different populations: bacteria that actively proliferate during the first 24 hrs after invasion, leading to their release in extracellular milieu as a consequence of host cell death, and bacteria that do not replicate. These latter staphylococci can persist within host-cell cytoplasm for several days [3-9, 36]. Here we addressed the role of TKT in these two phenotypes in EA.hy926 human endothelial cells by using: i) time lapse-microscopy, to evaluate early intracellular proliferation of *S. aureus* and ii) colony forming units counting, to monitor the survival kinetics of internalized bacteria over a ten days period.

#### Time-lapse video microscopy

EA.hy926 cells expressing mKate2 nuclear fluorescent protein were infected by GFP-expressing bacteria. Gentamicin was used throughout the experiment to eliminate extracellular bacteria and prevent re-infection (see Methods), allowing us to solely focus on the fate of intracellular persistent bacteria. Cell infection was monitored using an IncuCyte^®^ S3 imaging system and images were captured every 30 minutes. Bacterial entry into endothelial cells was similar for WT and ∆*tkt* strains, showing 0.3~0.5 GFP expressing particles recorded per cell (**Figure 2A-B**). Active proliferation of WT *S. aureus* inside EA.hy926 cells was confirmed after 24 hours of infection (**Figure 2A-B and Video 1)** with an average number of GFP expressing particles rising from 0.5 to 2 per cells. At the single cell level, WT *S. aureus* showed active intracellular multiplication, ultimately leading to cell death and bacterial release in the extracellular medium (**Figure 2A-B and Video 1**). Thanks to the permanent presence of gentamicin in the medium bacterial release did not lead to proliferation in the extracellular medium. In contrast, the ∆*tkt* mutant strain showed limited proliferation during the first 24 hours (**Figure 2A-B and Video 2**). Finally, using LAMP-1 colocalization assay (Lysosome-associated membrane proteins 1), we confirmed that the intracellular ∆*tkt* strain growth defect was not due to a decreased phagosomal escape (**Supplementary Figure S4**).

**Figure 2.**
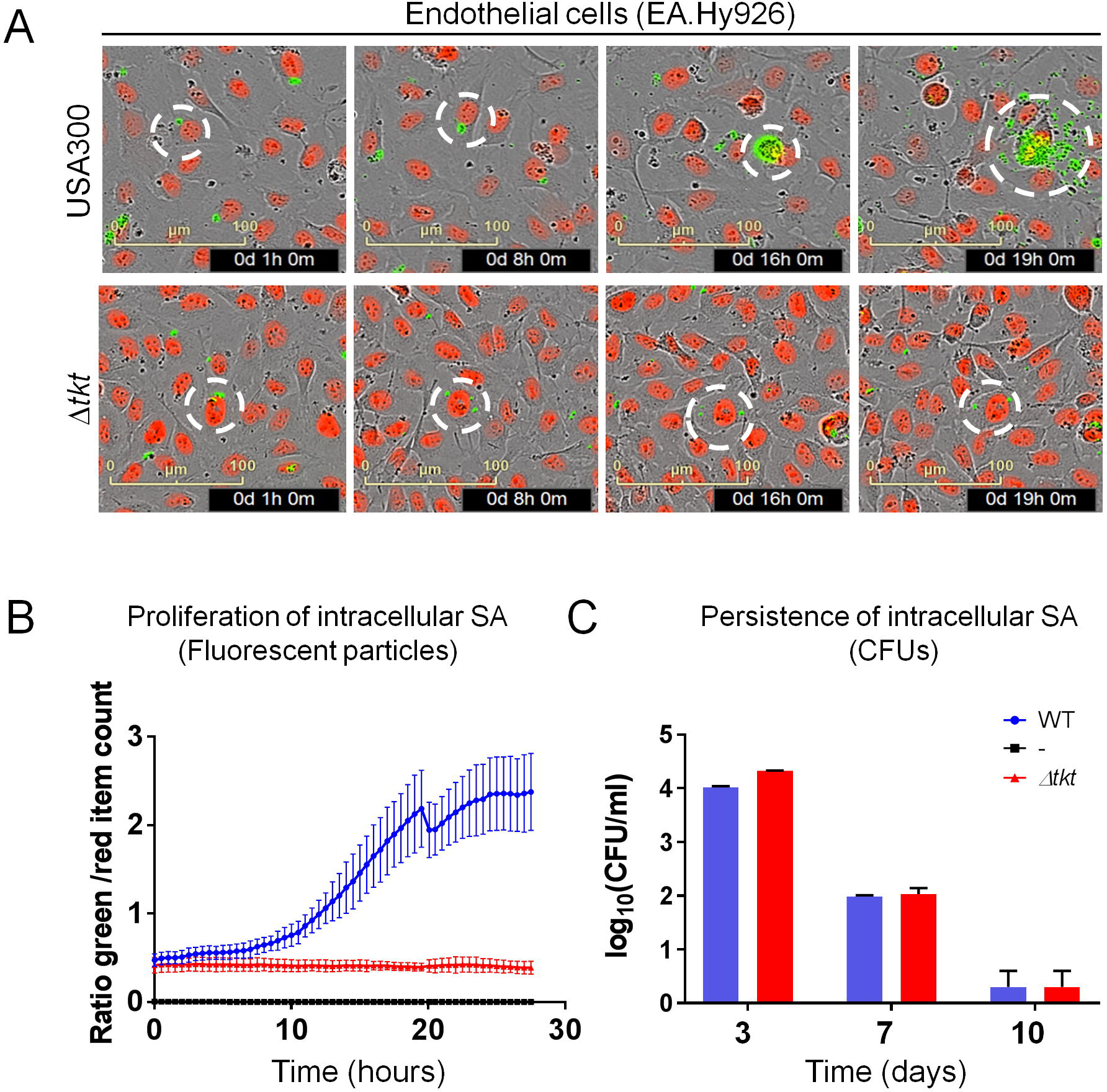
Intracellular proliferation and persistence of *S. aureus*. **A, B. Intracellular proliferation of USA300 wild type (WT) and Δ*tkt* mutant (Δ*tkt*) strains.** Endothelial EA.Hy926 cells expressing mKate2 nuclear restricted red fluorescent protein were infected with GFP expressing WT and Δ*tkt* strains. One hour after infection, cells were washed several times with gentamicin containing medium to eliminate extracellular bacteria. Gentamicin concentration of 50µg/ml was maintained throughout the experiment. Images were acquired every 30 minutes using Incucyte^®^ S3 live cell imaging system. A. Representative images (see also video S1 and S2). Red: cell nuclei. Green: GFP expressing bacteria. For each line, a white circle was centred on a unique infected cell. Bar = 100µm. B. Quantitative analysis representing the number of green particles divided by the number of red nuclei during the first day after infection. Blue line: WT strain. Red line: Δ*tkt* strain. Black line: non infected cells. **C. Persistence of intracellular WT and Δ*tkt* strains.** Endothelial EA.Hy926 cells were infected with WT and Δ*tkt* strains as described above. Gentamicin was maintained throughout the experiment. Survival of *S. aureus* was assessed by plating and counting CFUs three, seven and ten days after infection. Results are shown as mean ± SD of triplicate measurements.

#### Long-term intracellular survival

Using the same infection conditions as for time-lapse experiment, we monitored survival of internalised Δ*tkt* mutant compared to that of the WT strain at three, seven or ten days after infection by CFU counting. The Δ*tkt* strain behaved likes the WT strain throughout the infection period, ultimately leading to its complete elimination after 10 days (**Figure 2C**).

Thus, the role of TKT during *S. aureus* host cell infection seems to be essential for bacteria proliferation and restricted to early time points.

### *tkt* inactivation impairs proliferation of *S. aureus* in the kidney of infected mice

The ability of internalized *S. aureus* to readily proliferate inside host cell cytoplasm may be seen as an effective mechanism to avoid host innate immunity, ultimately allowing an increased colonization of the infected site. In this respect, it has been previously shown in the mouse model that after intravenous (IV) injection, although rapidly cleared from the bloodstream, bacteria were able to proliferate in the kidney, leading to abscesses formation within the first days of infection [37].

We monitored the kinetics of *in vivo* proliferation of WT and Δ*tkt* strains in kidneys and spleens of BalbC mice (**Figure 3**) after IV infection with 5×10^6^ bacteria per mouse. The bacterial burden in kidneys and spleens was quantified by plating CFUs at 4 h, 24 h, 48 h and 72 h after infection. We observed no significant differences between the bacterial counts in the spleens of mice infected either with WT or the Δ *tkt* mutant. In both cases, a progressive reduction of the bacterial burden was recorded (ranging from 3.6×10^5^ and 1.3×10^5^ at 4 h; down to 2.8×10^3^ and 0.9×10^3^ at 72 h, for WT and ∆*tkt* respectively). In contrast, in kidneys, whereas the WT strain readily proliferated (from 6.4×10^4^ ± 5.10^3^ WT *S. aureus* per organ at 4 hrs to 1.10^8^ ± 6.10^7^ WT *S. aureus* per organ at 72 hrs), multiplication of the Δ *tkt* mutant was severely impaired at all time-points tested, with a *ca.* 10^5^-fold reduction of ∆*tkt* counts at 24h compared to WT counts. This result confirms the importance of TKT -and most likely of the PPP-in bacterial proliferation *in vivo* and especially during the first 24 hours after infection.

**Figure 3.**
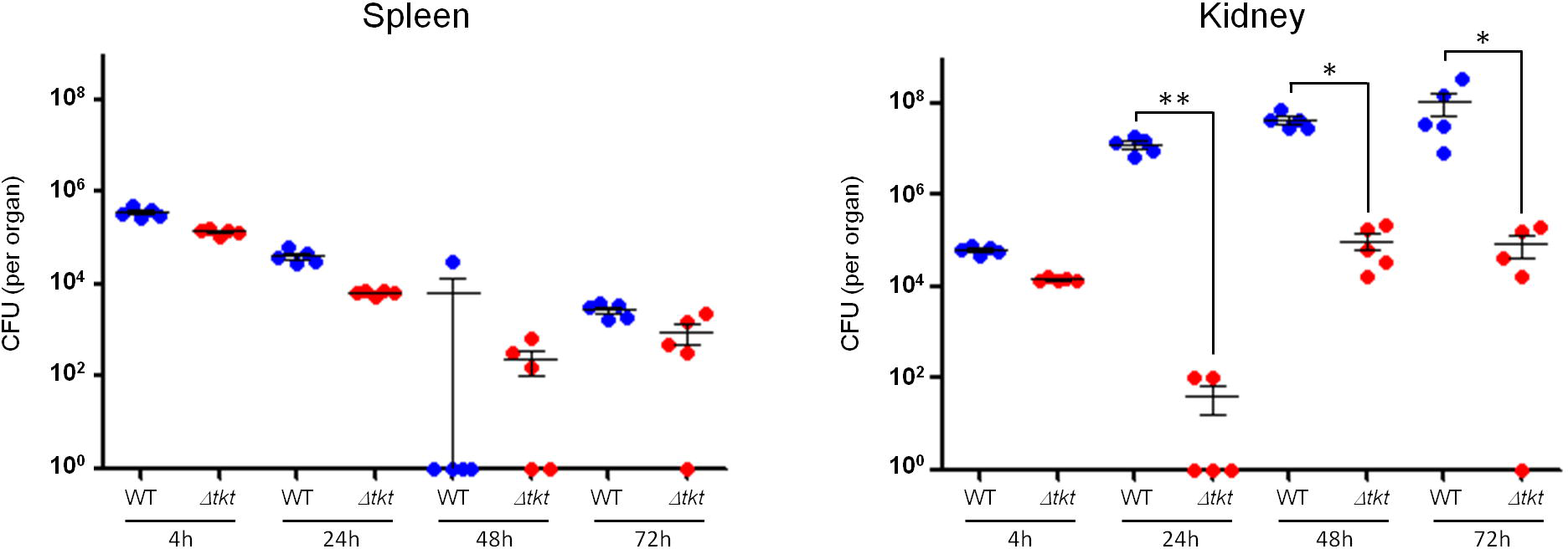
Proliferation of *S. aureus* in the spleen and the kidney of infected animals. Bacterial loads in the spleen and the kidney of BalbC mice infected intravenously with 5.10^6^ bacteria of the WT and Δ*tkt* strain. Bacterial counts are expressed in CFU/organ (n = 5). Data are shown as mean ± SD. Statistical significance was determined using the Kruskal Wallis test (* *p*< 0.05, ** *p*<0.01).

### *tkt* inactivation is associated with and altered *rpiRc* and *sigB* expression

*S. aureus* is known to sense its environment and to regulate virulence factors depending on the availability of carbon sources. For example, recent studies have shown that the transcriptional regulator RpiRc could sense metabolic shifts, especially in the PPP, and control expression of RNAIII [29, 30]. RNAIII is a key effector in the quorum-sensing system that regulates the expression of a large number of virulence factors [38]. RpiRc is also linked to the alternative Sigma B factor (SigB) that has pleiotropic roles in gene regulation and is a master regulator of intracellular survival [29, 39, 40].

These data prompted us to follow the expression profile of the master regulator *sigB* and *rpiRc* in the WT and the Δ*tkt* mutant strains during endothelial cells invasion (**Figure 4**). EA.hy296 cells were infected as described above to solely focus on intracellular bacteria. Between day 1 and day 7, *sigB* and *rpiRc* transcription increased in the WT strain (by 4-fold and 3-fold, respectively) (**Figure 4**). In sharp contrast, as early as one day after infection, both genes were significantly down-regulated in the ∆*tkt* mutant strain compared to WT (8-fold and 7-fold decrease for *sigB* and *rpirC* expression, respectively). Consistent with reports demonstrating inhibition of RNAIII expression by RpirC and SigB [18], we observed a dramatic increase in the expression of *RNAIII* in the ∆*tkt* mutant strain (**Figure 4**), suggesting that the Δ*tkt* mutant strain is unable to control *rpiRc* and *sigB* expression during intracellular survival. To further confirm these data, we monitored the expression of two major regulators of SigB activity, RsbU and RsbW [41]. RsbW, which is co-expressed with SigB, is known to sequestrate and inhibit SigB. RsbU, whose expression is under the control of SigA, is a phosphatase involved in the dephosphorylation of RsbV. In turn, dephosphorylated RsbV (dRsbV) is able to sequestrate RsbW, allowing the release of active SigB. RsbU is thus considered as an activator of SigB, allowing a fine-tuning of SigB activity.

**Figure 4.**
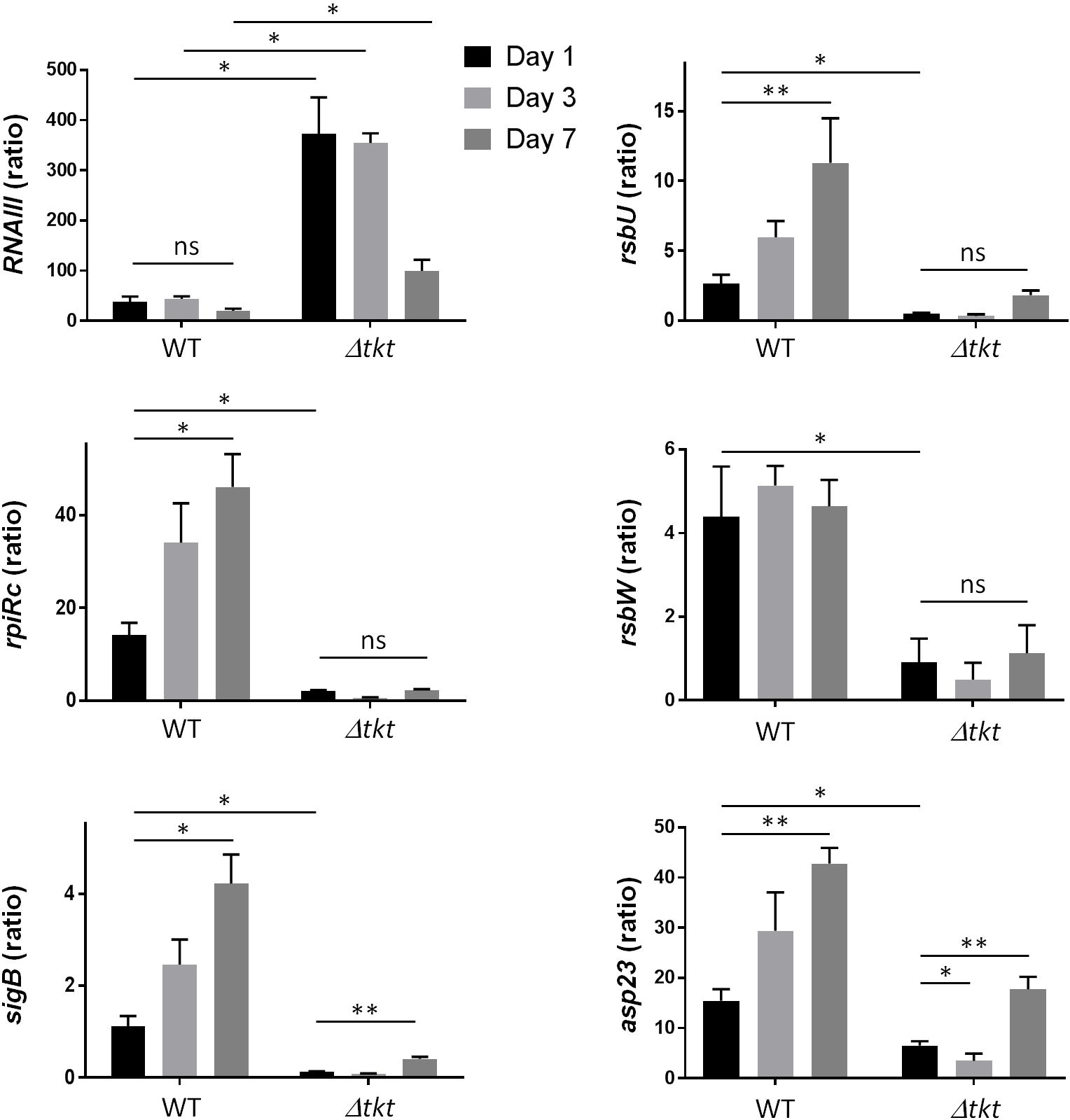
Transcription profile of intracellular *S. aureus*. Endothelial EA.Hy926 cells were infected with USA300 WT and Δ*tkt* strain as described in figure 2. One, three or seven days after infection cells were washed and collected. RNA from intracellular *S. aureus* were prepared and gene expression of *RNAIII*, *rpiRc*, *sigB*, *rsbU, rsbW* and *asp23* was analysed by quantitative RT-PCR. Gene expression level was normalized by that of *gyrA* at days 1, 3 and 7 after infection of EA-hy296 cells. Data are shown as mean ± SD of triplicate measurements from two independent experiments. (ANOVA, n.s. : not statistically different; * *p*<0.01, ** *p*<0.001).

We found that the transcription of both *rsbW* and *rsbU* genes was down-regulated in the Δ*tkt* mutant during infection of endothelial cells. The down-regulation of *rsbW* is consistent with the down-regulation of its co-expressed gene *sigB* (**Figure 4**). The concomitant down-regulation of *rsbU* suggests a global inhibition of the *sigB* pathway in the Δ*tkt* mutant strain. Finally, we followed expression of *asp23,* a *sigB*-dependent locus [42]. The recorded down-regulation of *asp23* transcription in the Δ*tkt* mutant strain confirmed the inhibition of SigB-dependent transcription. Altogether these transcriptional analyses suggest that the metabolic dysregulation observed in Δ*tkt* strain is sufficient to promote a dramatic reprogramming of the RpiRc and SigB pathways when bacteria invade human cells.

### The SigB pathway may be controlled by modulating carbon source availability

Inside the cytoplasm of host cells the Δ*tkt* mutant is likely to deal simultaneously with PPP blocking and altered levels of carbon-based nutrients compared to the outside compartment. The above results suggested that the PPP might take control of *rpiRc* and *sigB* expression when bacteria invade endothelial cells. We thus hypothesized that PPP-associated carbon sources may possibly influence *rpiRc* and/or *sigB* expression *in vitro*. We therefore grew WT USA300 in CDM supplemented either with glucose (CDM-glucose), ribose (CDM-ribose) or fructose (CDM-fructose) a carbon source possibly fuelling the PPP) and quantified by qRT-PCR transcription of *sigB*, *rpiRc*, *rsbU/W* and *asp23* genes. Remarkably, we found that the regulation of the whole *sigB* operon was impacted by the carbon source used in CDM (**Figure 5**). Indeed, expression of *sigB, rsbU/W* and *rpiRc* decreased in CDM-fructose or CDM-ribose, compared to CDM-glucose. In contrast, *asp23* expression was only decrease by 2-fold in CDM-fructose and increased by 3-fold in CDM-ribose, suggesting that in this growth conditions, *asp23* expression might be regulated by other factor than *sigB* (**Figure 5**). Hence, the availability of carbon sources, and especially those related to PPP, may be sensed by the bacteria and affect the regulation of *rpiRc* and *sigB* expression.

**Figure 5.**
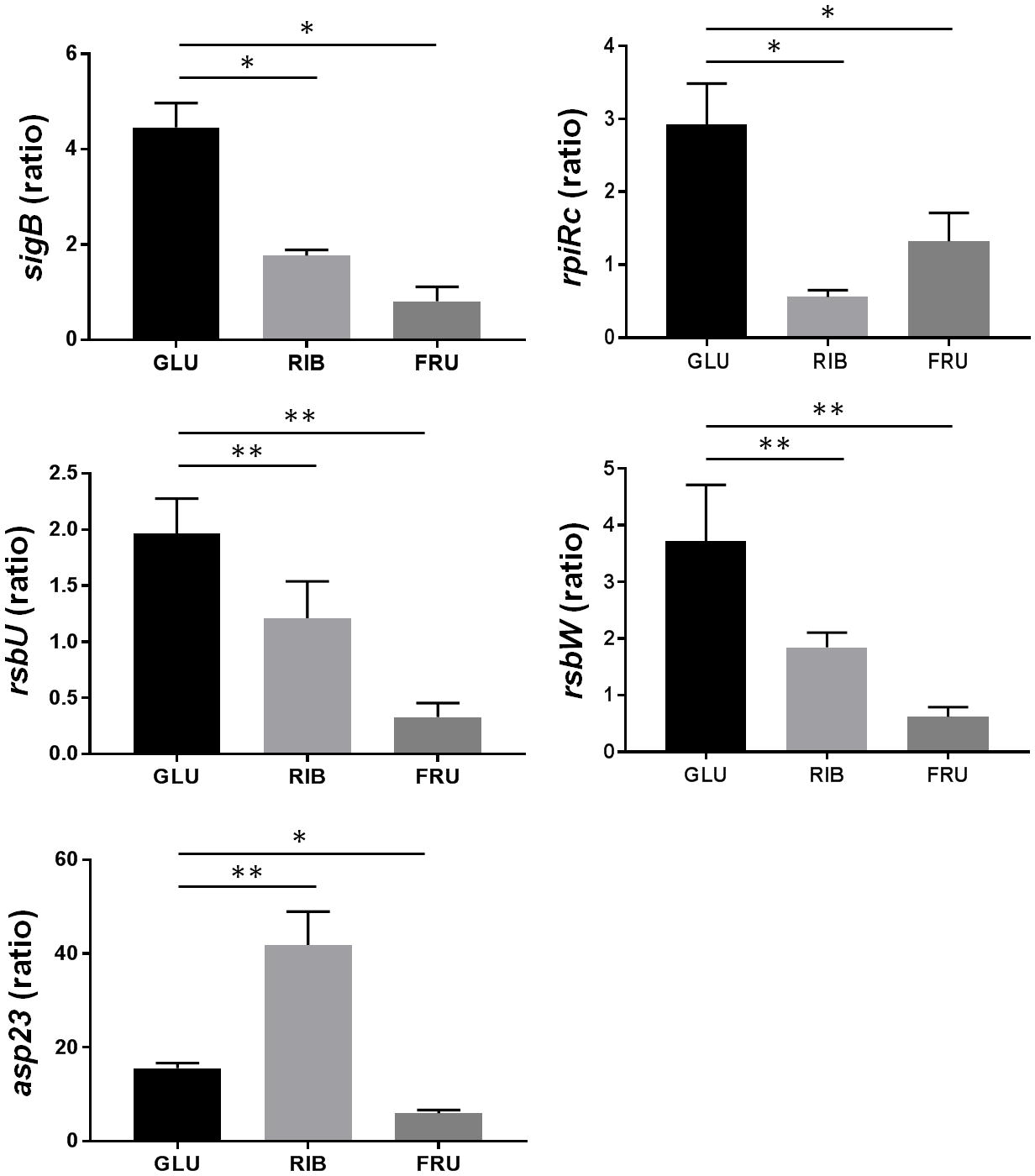
Carbon sources influence the transcription profile of the SigB pathway. *S. aureus* USA300 WT strain was grown to stationary phase in CDM supplemented with glucose or ribose or fructose and gene expression profile of *rpiRc*, *sigB*, *rsbU, rsbW* and *asp23* was analysed by quantitative RT-PCR. Gene expression level was normalized by that of *gyrA*. Data are shown as mean ± SD of triplicate measurements from two independent experiments. Statistical significance was assessed in comparison to bacteria grown in CDM glucose (ANOVA, * *p*<0.01, ** *p*<0.001).

## Discussion

Inactivation of the *tkt* gene, encoding the unique transketolase of *S. aureus* USA300, led to a major decreased in early intracellular proliferation and impaired *in vivo* multiplication in a kidney colonization model. Our results suggest that the intracellular proliferation defect of the ∆*tkt* mutant occurs through the inhibition of SigB and RpiRc expression and the concomitant increased RNAIII transcription, thus highlighting a novel connection between the PPP and these master regulators during cell invasion.

Our whole cell metabolomic analysis of planktonic-grown bacteria (**Figure 1, Supplementary Figure S2 and Table S1**) showed a huge increase in R5P intermediates in the ∆*tkt* mutant compared to WT that could be explained by the impaired entry of R5P into glycolysis and a concomitant significant decrease in SH7P relative concentration. In *S. aureus*, ribose is imported by the RbsU transporter and converted to R5P by the RbsD/RbsK pathway. RbsR, the repressor of *rbsUDK* operon, is itself highly regulated by SigB [43]. Interestingly, *S. aureus* grown in CDM-ribose showed a decreased expression of *sigB*. Hence, stress signals that modulate SigB activity are likely to be important effectors for controlling the quantity of RbsR, thereby affecting ribose uptake.

Significant changes in the relative concentrations of TCA cycle intermediates were also recorded in the ∆*tkt* mutant compared to WT (**Supplementary Figure S3**). The TCA cycle plays a central role in maintaining the bacterial metabolic status and has been repeatedly implicated in the regulation of staphylococcal virulence [44, 45]. Hence, changes in TCA cycle activity are likely to induce alterations of the overall metabolome of the bacterium that, in turn, may modulate the activity of metabolite-responsive global regulators such as SigB or RpiR.

RpiRc, a RpiR family-member controlling enzymes involved in sugar catabolism in *S. aureus*, has been shown to repress RNAIII [28], supporting the notion of a direct connection between the PPP and the regulation of *S. aureus* virulence. The expression of SigB has also been shown to play a crucial role during the intracellular life of bacteria possibly by down-regulating pro-inflammatory virulence factors and increasing the expression of factors promoting persistence [18, 40].

We propose (**Figure 6**) that in wild-type *S. aureus*, transketolase activity results in a positive regulation of *rpiRc* and *sigB* transcription and allows the repression of RNAIII. Consequently, repression of toxins production favours intracellular growth. In contrast, in a ∆*tkt* mutant, *rpiRc* and *sigB* transcriptions are not activated, possibly due to metabolic changes (such as R5P accumulation) and RNAIII is not repressed. At this stage, it cannot be excluded that the enzyme itself might also directly affect transcription of these regulators.

**Figure 6.**
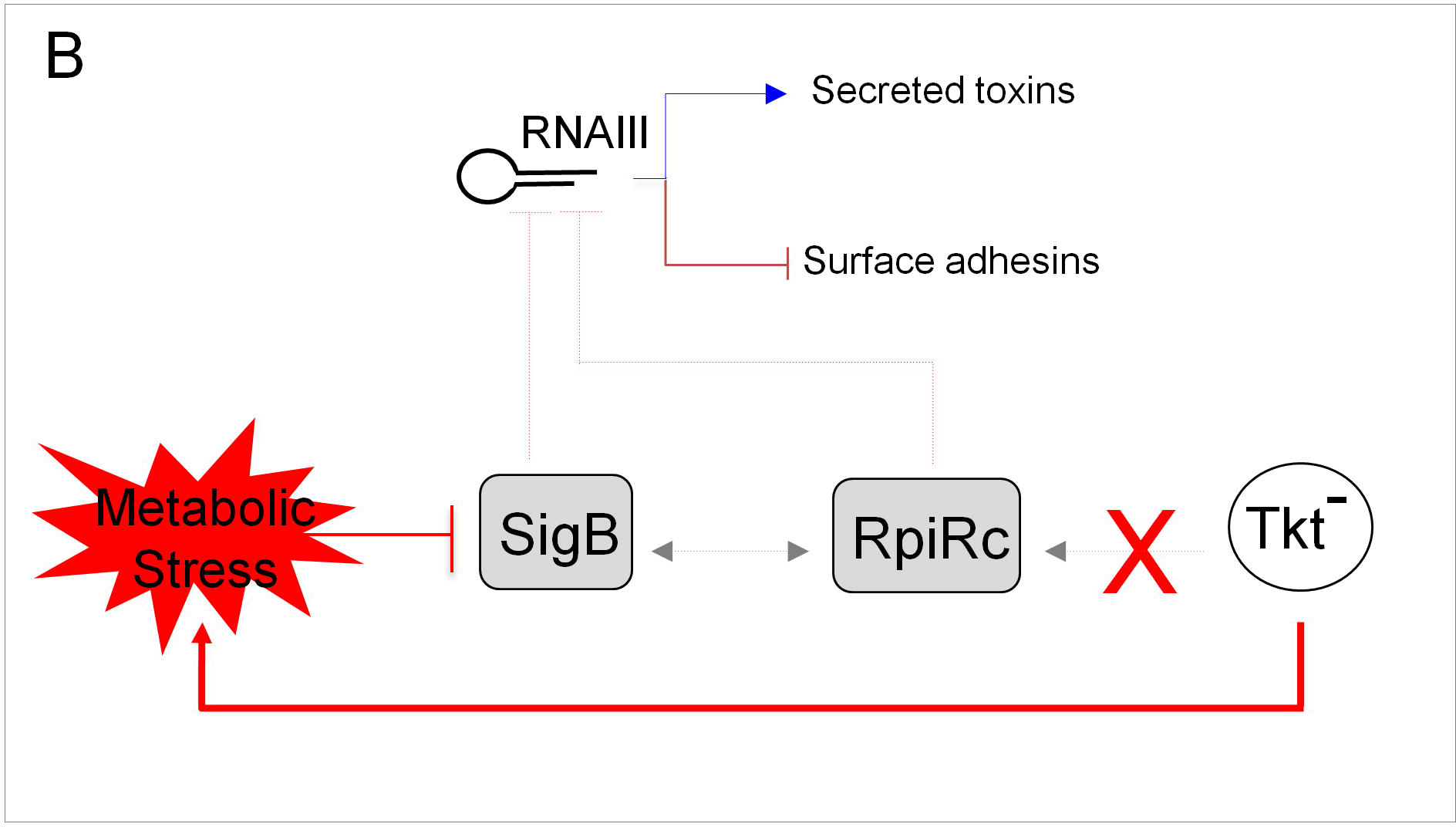
Schematic depiction of the impact of TKT inactivation on *sigB* and *rpiRc* regulation. SigB dependent regulation of *RNAIII* in *Δtkt* context.

TKT inhibitors are currently actively tested in cancer therapy [46] and TKT could constitute an efficient target against tuberculosis [35] and malaria [47]. The present work, highlighting an unprecedentedly reported role of transketolase in *S. aureus* intracellular survival, suggests that modulation of the pentose phosphate pathway activity may also represent an interesting mean to fight *S. aureus* infections.

---

**Movie 1: Intracellular proliferation of USA300 wild type strain.** Endothelial EA.Hy926 cells expressing a nuclear restricted Red Fluorescent Protein were infected with GFP expressing wild type strains. One hour after infection cells were washed several times with gentamicin containing medium to eliminate extracellular bacteria (see materials and methods section). Gentamicin concentration of 50µg/ml was maintained throughout the experiment. Images were acquired every 30 minutes using Incucyte^®^ S3 live cell imaging system.

**Movie 2: Intracellular proliferation of USA300 Δ*tkt* mutant strain.** Same protocol as above in movie S1.

## Acknowledgements

We are grateful to the Cell imaging core facility of the “Structure Fédérative de Recherche” Necker INSERM US24/CNRS UMS3633 for his technical support. The following reagents were provided by the Network on Antimicrobial Resistance in *Staphylococcus aureus* (NARSA) for distribution by BEI Resources, NIAID, NIH: *Staphylococcus aureus*, Strain USA300-0114, NR-46070; *Escherichia coli* – *Staphylococcus aureus* Shuttle Vector pNR-46158, Recombinant in *Staphylococcus aureus*, NR-46158.

## Notes

Conflict of interest: No potential conflict of interest was reported by the authors.

**Funding**: These studies were supported by INSERM, CNRS and Université Paris Descartes Paris Cité Sorbonne. Xin Tan was funded by a scholarship from the China Scholarship Council (n° CSC NO. 201508500097). The funders had no role in study design, data collection and analysis, decision to publish, or preparation of the manuscript.

