## Supplementary figures and images for "Transketolase of *Staphylococcus aureus* is involved in the control of master regulators of stress response during infection"

### Supplementary figure S1

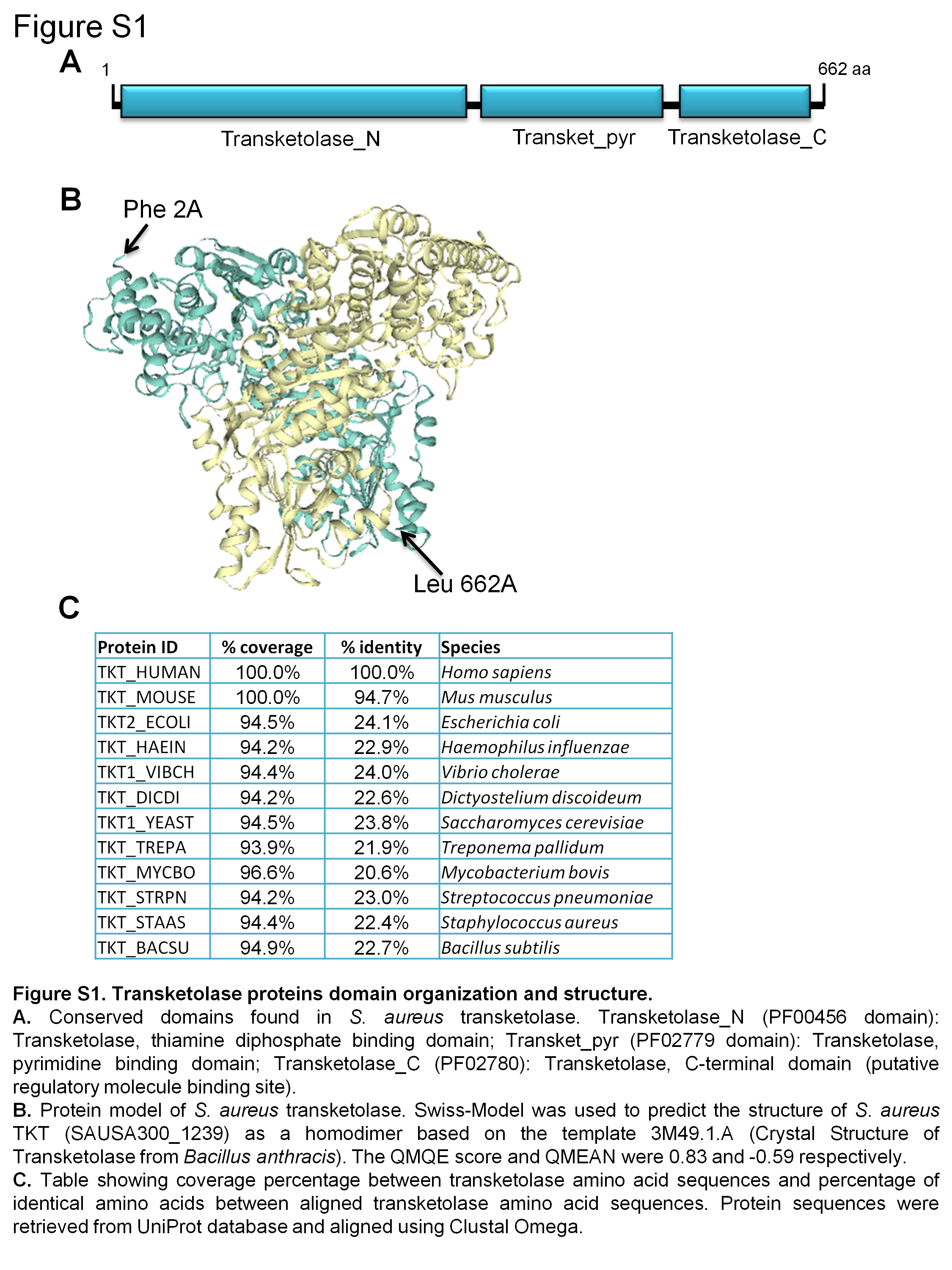

### Supplementary figure S2

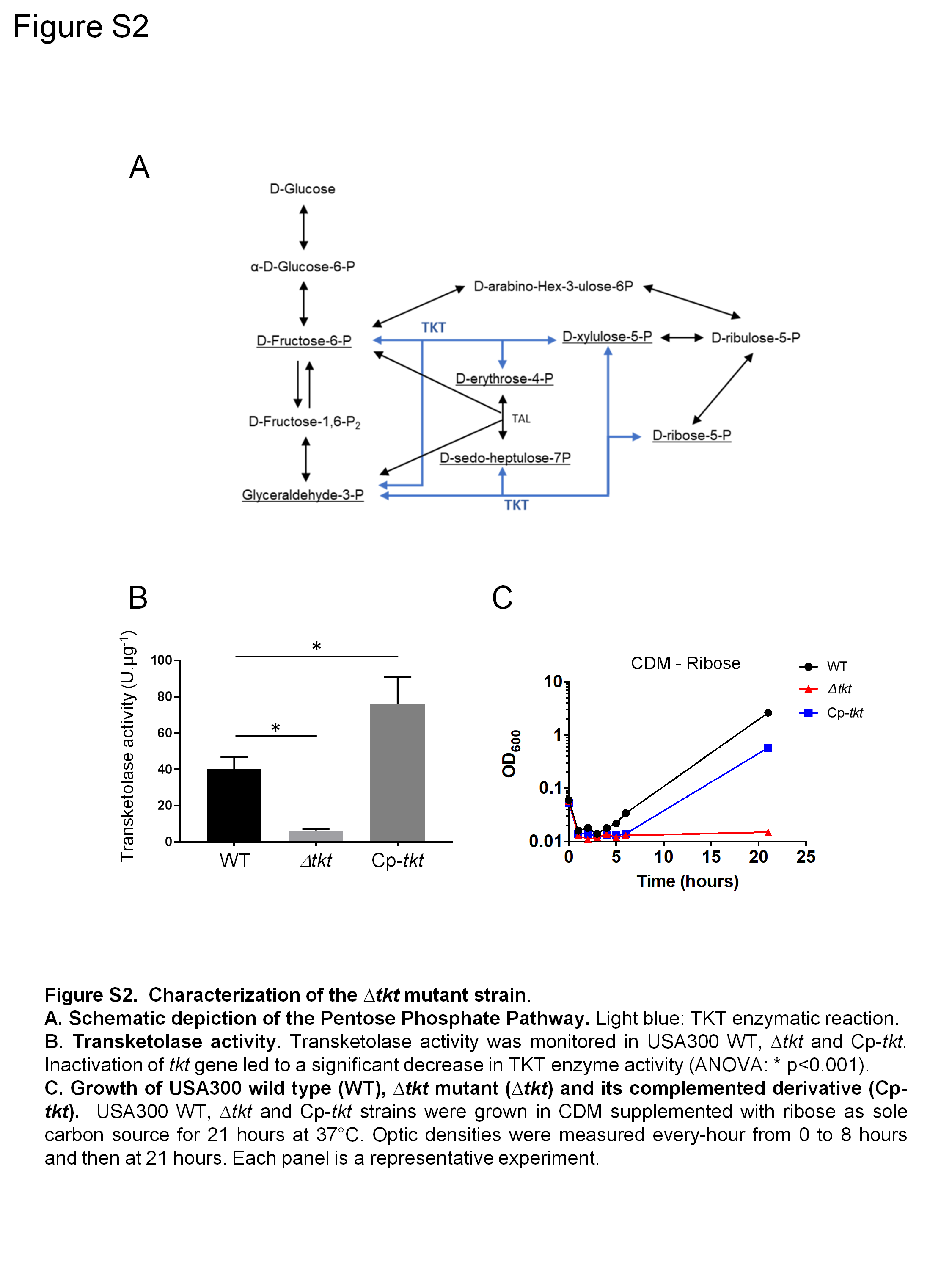

### Supplementary figure S3

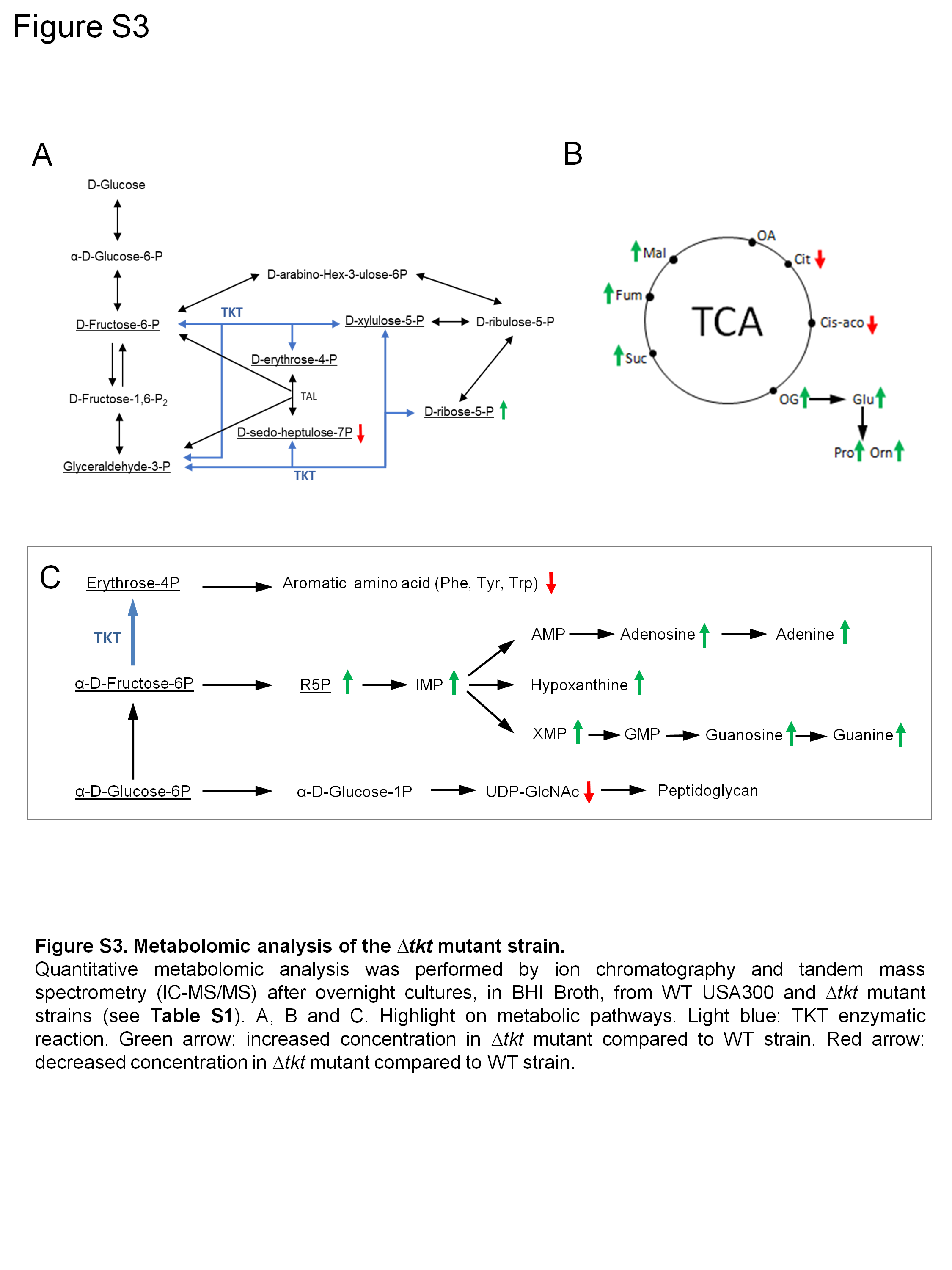

### Supplementary figure S4

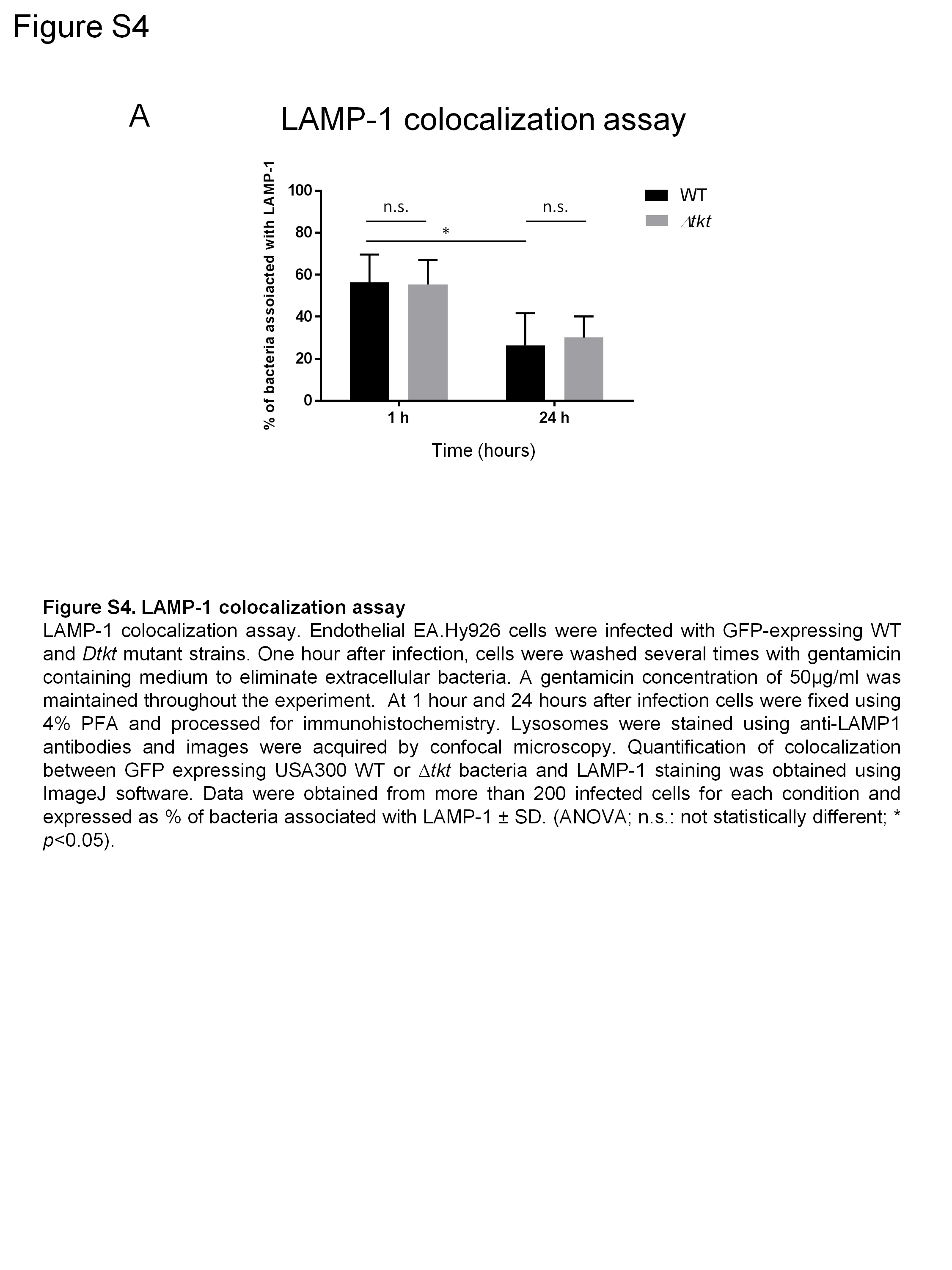
